# Mutations within the predicted fragment-binding region of FAM83G/SACK1G abolish its interaction with the Ser/Thr kinase CK1α

**DOI:** 10.1101/2025.06.25.661481

**Authors:** Javier S. Utgés, Diane Lee Zhi Xuan, Brune Le Chatelier, Lorraine Glennie, Thomas J. Macartney, Nicola T. Wood, Geoffrey J. Barton, Gopal P. Sapkota

## Abstract

SACK1G (aka FAM83G, PAWS1) plays a central role in activating canonical WNT signalling via interaction with the Ser/Thr kinase CK1α. This loss of CK1α binding and WNT signalling underlies the pathogenesis of Palmoplantar Keratoderma (PPK) caused by several reported mutations in the *SACK1G* gene. We modelled the scaffold anchor of CK1 (SACK1) domain of SACK1G and used fragment-bound structures of the SACK1B (FAM83B) dimer to guide our analysis. This allowed us to computationally predict several key residues near the fragment-binding site in SACK1G that may be important for its function. We mutated these residues, introduced them into *SACK1G*^-/-^ DLD-1 colorectal cancer cells and investigated their ability to bind endogenous CK1α. We uncovered two SACK1G mutations, namely Y204A and I206A, that abolish interaction with CK1α similarly to the PPK pathogenic mutant A34E. Consistent with this loss of SACK1G-CK1α interaction, the molecular glue degrader of CK1α, DEG-77, fails to co-degrade the Y204A and I206A mutants while it still co-degrades native SACK1G. Our findings demonstrate the utility of our computational methods to uncover functional residues on proteins based on fragment-binding sites.

## Introduction

The eight members (A-H) of the FAM83 family of proteins are characterised by the presence of the conserved <u>S</u>caffold <u>A</u>nchor of <u>CK1</u> domain (SACK1; formerly known as the Domain of Unknown Function DUF1669) but are otherwise different in sequence composition and length ^1,2^. The CK1 family of Ser/Thr protein kinases control a myriad of cellular processes, including cell cycle progression, circadian rhythms, WNT signalling and Sonic Hedgehog signalling ^3^. Contrary to the dogma that CK1 isoforms are constitutively active kinases, their activity, subcellular localisation, and stability in cells is tightly regulated to allow phosphorylation of specific substrates and effectuate distinct biological outcomes, although how this is achieved is poorly understood ^3^. In a series of recent discoveries, we identified the eight members of the FAM83/SACK1 family of proteins (hereafter referred to as SACK1 domain-containing proteins A-H, or SACK1A-H) as master regulators of CK1 isoforms ^2,4–9^. The conserved SACK1 domain of SACK1A-H proteins mediates their interaction with CK1 isoforms and dictates the subcellular distribution – and potentially substrates – of the interacting CK1 isoforms ^2^. The precise molecular and biological roles of most SACK1 proteins are poorly defined. Among the best characterised functionally, the SACK1F-CK1α and SACK1G-CK1α interactions are known to drive the activation of canonical WNT signalling ^4,8^, while SACK1D-CK1α interaction at the mitotic spindle has been reported to ensure proper spindle orientation and timely and error-free progression through cell division cycle ^10^.

Four different missense mutations, resulting in A34E, R52P, F58I and R265P substitutions, within the SACK1 domain of SACK1G have been reported to cause Palmoplantar Keratoderma (PPK), which is characterised by thickening of the skin on the palms and soles of the feet and abnormal hair growth, in both humans ^5,9,11,12^ and dogs ^13^. These mutations are reported to abolish SACK1G-CK1α binding ^5,9,12^ and attenuate canonical WNT signalling ^5,9^, highlighting the importance of CK1α binding for SACK1G function. Interestingly, a spontaneous deletion of a large part of the *SACK1G* gene from mice was reported to cause ‘*wooly*’ hair phenotype ^14^. The functional convergence between SACK1G and CK1α is also highlighted by the similar PPK phenotypes evident in human patients displaying a loss in *SACK1G* expression ^12^ and transgenic mice targeted for ablation of CK1α expression from keratinocytes ^15^.

The crystal structures of the SACK1 domains of FAM83A/SACK1A (PDB: 4URJ ^16^) and FAM83B/SACK1B (PDB: 5LZK ^17^) have been solved. The structures reveal a unique dimer with PLD1-like fold ^18^. However, the prototypic catalytic “HKD” triad of PLD1 is not conserved in SACK1 proteins and no PLD activity for any of the SACK1 proteins has been reported. A structure-based fragment screening undertaken using the SACK1 domain of FAM83B/SACK1B has reported ten unique fragments bound to the SACK1 domain across eleven structures (PDB: 5QHI ^19^, 5QHJ ^20^, 5QHK ^21^, 5QHL ^22^, 5QHM ^23^, 5QHN ^24^, 5QHO ^25^, 5QHP ^26^, 5QHQ ^27^, 5QHR ^28^ and 5QHS ^29^). No structures of SACK1 domains from other SACK1 proteins have been reported nor are there reports of any known binding fragments or ligands. In this work, AlphaFold Multimer version 3 ^30–32^ was employed to generate 3D structure models of a SACK1G dimer. Using a combination of solvent accessibility, evolutionary divergence and enrichment in human population variation ^33–36^, residues within a conserved binding site between SACK1B and SACK1G were ranked by their likelihood of function and prioritised for experimental validation.

## Methods

### Computational Methods

#### Structure prediction

Protein three-dimensional (3D) structure models were generated using the Colabfold version 1.5.2 implementation ^37,38^ of AlphaFold Multimer (AFM) v3 ^30–32^. Additionally, secondary structure predictions were obtained from the JPred 4 web server ^39–42^. Protein sequences were obtained from UniProt ^43^ and domain annotations extracted from InterPro ^44,45^. Models were obtained for dimers of the scaffolding anchor of CK1 (SACK1) domain (PF07894) – previously known as DUF1669 – of human SACK1B (Q5T0W9) and SACK1G (A6ND36). Sequence constructs included residues 1-286 and 1-315 for SACK1B and SACK1G, respectively, to ensure the full SACK1 domain was modelled. Models were visualised with ChimeraX ^46–48^. Model quality was assessed at the interface level with the interface predicted template modelling (ipTM) score as well as the new interaction prediction score from aligned errors (ipSAE) ^49^ with *dist* and *PAE* thresholds of 15 Å. Predicted local distance difference test (pLDDT) and predicted aligned error (PAE) were utilised to measure model quality at the residue level.

#### Fragment screening analysis

Eleven structures from a fragment screening experiment on the SACK1 (DUF1669) domain dimer of human SACK1B (Q5T0W9) were extracted from the PDBe ^50^. Twenty-four ligand molecules, ten of which were unique, binding to the SACK1B dimer across the eleven structures, were grouped into six distinct ligand binding sites by the method of Utgés *et al.* ^35,51,52^. Binding sites and their residues were characterised by evolutionary divergence across homologues and human population missense variation constraint. jackHMMER ^53^ was employed to build a multiple sequence alignment for FAM83B and evolutionary divergence was quantified with the normalised Shenkin ^54^ divergence score, *N_Shenkin_* ^34^. Residue-level constraint within the human population was measured with a missense enrichment score (MES) ^33,36,55^ and variants from gnomAD ^56^.

*N_Shenkin_* is a divergence score ranging from 0 to 100. Positions that are conserved across homologues present lower divergence scores, *N_Shenkin_* < 25, while divergent positions present higher scores, *N_Shenkin_* > 75. MES is calculated as the natural logarithm of an odds ratio (OR) and expresses the likelihood of observing missense variants in an alignment column, or residue position, relative to the rest of the alignment (or protein). Positions enriched in missense variation present MES > 0. Positions that are depleted in missense variation, i.e., are constrained within human, present MES < 0. Neutral positions that are neither enriched nor depleted present MES = 0. Refer to our previous work ^33–36^ for more details on how these scores are calculated and interpreted.

### Biological Methods

#### Materials

Anti-CK1α (SA527), anti-SACK1G (S685C), and anti-SACK1F (SA103) antibodies and Protein-G Sepharose beads coupled to anti-GFP nanobody were generated by the MRC PPU Reagents and Services. Anti-GAPDH antibody (Cat #2118) and goat anti-Rabbit IgG (H+L)-HRP (#7074) were from Cell Signaling Technology. Rabbit anti-sheep IgG (H+L)-HRP (#31480), Gibco DMEM and Fetal Bovine Serum (FBS) were obtained from Thermo Fisher Scientific. All plasmids employed in this study were generated by MRC PPU Reagents and Services at University of Dundee, with detailed information accessible at https://mrcppureagents.dundee.ac.uk. DEG-77 was a kind gift from Christina Woo (Harvard).

#### Cell culture

Human embryonic kidney cells 293FT, wild type DLD-1 colorectal cancer cells and *SACK1G ^-/-^* DLD-1 cells ^8^ were cultured in Dulbecco’s Modified Eagle’s Medium (DMEM) supplemented with 10% FBS, 2 mM L-glutamine and 1% streptomycin/penicillin, maintained at 37°C in 5% CO_2_ in humidified incubators. Indicated concentrations of DEG-77 were diluted in DMSO and added to cells, mixed by swirling and returned to the incubators for the indicated length of time prior to lysis. For production of retroviral vectors, the following were cloned into pBabeD-puromycin plasmids: WT SACK1G-GFP (DU20988), SACK1G^A34E^-GFP (DU29572) SACK1G^V175A^-GFP (DU71867), SACK1G^I206A^-GFP (DU71868), SACK1G^Q301A^-GFP (DU71869), SACK1G^F260A^-GFP (DU71874), SACK1G^Y204A^-GFP (DU71875), and SACK1G^R235A^-GFP (DU71885). All constructs were sequence-verified by the DNA Sequencing Service, University of Dundee (http://www.dnaseq.co.uk). All constructs are available to request from the MRC-PPU Reagents and Services webpage (http://mrcppureagents.dundee.ac.uk) and the unique identifier (DU numbers) provide direct links to the cloning strategies and sequence details.

#### Retroviral generation of stable cell lines

*SACK1G^-/-^* DLD-1 cells stably expressing the different SACK1G-GFP mutants were generated via retroviral transduction by following established biological safety guidelines. Briefly, the pBabeD-puromycin vectors encoding the specific mutants (6 μg), pCMV5-gag/pol (3.2 μg) and pCMV5-VSV-G plasmids (2.8 μg) (Clontech) were co-transfected into ∼60-70% confluent HEK-293FT cells in a 10-cm diameter dish using PEI reagent as previously described ^57–59^. The medium was replaced with fresh medium after 16 h and the medium containing the retroviral particles was collected 48 h post-transfection, filtered through a 0.45 mm filter, and added to target *SACK1G ^-/-^* DLD-1 cells along with 10 μg/ml polybrene (Sigma-Aldrich). 48 h later, successfully transduced cells were selected in fresh medium containing 2 μg/ml puromycin (Sigma-Aldrich) for a further 48 h.

#### Cell Lysis

The cells were washed twice with ice-cold PBS and scraped on ice in lysis buffer (LB: 50 mM Tris-HCl (pH 7.5), 270 mM sucrose, 150 mM sodium chloride, 1 mM EDTA (pH 8.0), 1 mM EGTA (pH 8.0), 1 mM sodium orthovanadate, 10 mM sodium β-glycerophosphate, 50 mM sodium fluoride, 5 mM sodium pyrophosphate, 1% (v/v) NP-40) supplemented with 1x cOmplete™ protease inhibitor cocktail (Roche) and 1 mM DTT. Cell lysates were transferred to Eppendorf tubes and vortexed briefly. The cell suspension was incubated for 15 min on ice to ensure complete lysis. The lysates were centrifuged at 17000 x g for 20 min at 4°C to remove cell debris, and cleared extracts were carefully transferred to new Eppendorf tubes. Protein concentration was measured using the Bradford assay and extracts were processed for SDS-PAGE & immunoblotting or immunoprecipitation (IP).

#### Immunoprecipitation

40 μL of 50% (v/v) slurry of anti-GFP-nanobody coupled Sepharose beads were added to cleared cell extracts (1 mg total protein) and IPs performed for 4 hours at 4°C on a rotating wheel. Pre-IP input samples and post-IP flowthrough (FT) extracts were collected to evaluate the IP efficiency and verify complete immune depletion of the target protein from the FT extracts. Following IP, the anti-GFP-nanobody coupled Sepharose beads were washed with 500 μL NP-40 lysis buffer thrice before IPs were resuspended in 50 μL of 1 x LDS sample buffer containing 1 mM DTT. The input and FT extracts were also reconstituted in 1 x LDS sample buffer containing 1 mM DTT and all samples were then denatured by boiling at 95 °C for 10 min prior to SDS-PAGE and immunoblotting.

#### SDS-PAGE and Immunoblotting

Reduced and denatured cell extracts containing equal amounts of protein (20 μg) and anti-GFP IP samples (50% of total) were resolved by SDS-PAGE and samples transferred to pre-equilibrated polyvinylidene fluoride (PVDF) membranes. Membranes were blocked in 5% (w/v) non-fat milk in TBS-T (20 mM Tris, 150 mM NaCl, 0.1% Tween-20) for 1 h at room temperature. Primary antibody diluted in 5% milk TBS-T was then incubated with the membranes overnight at 4°C on a shaking platform. Membranes were then washed 3x10 min in TBS-T with constant shaking and subsequently incubated with appropriate HRP-conjugated secondary antibodies (diluted in 5% milk/TBS-T), for 1 h at room temperature. Membranes were then washed 3x10 min in TBS-T and signal detection was performed by using chemiluminescence (Promega) on the ChemiDoc imaging system (Bio-Rad).

## Results

### SACK1B fragment screening analysis

Twenty-four fragments across eleven SACK1B structures were grouped into six different ligand binding sites (BS0 - BS5) using the method of Utgés *et al.* ^35^ (Figure 1 A). Binding sites 2, 3 and 5 present a Cluster 1 (C1) relative solvent accessibility label. Ligand sites in this cluster tend to be buried, conserved and depleted in missense variation in human. Moreover, C1 is strongly enriched in known functional sites ^35^. This suggests BS2, BS3 and BS5 might be functionally relevant. These three sites also present low sequence divergence across homologues and few missense variants (MES < 0) across human paralogues (Figure 1 B). This supports further their potential functional role. BS2 is at the predicted pseudo-PLD catalytic cleft, although whether it binds putative substrate phospholipids is yet to be determined ^1^ (Figure 1 C). BS3 is at the interface between the two SACK1B dimer copies (Figure 1 D) and BS5 is a buried pocket on the surface where twelve ligands bind (Figure 1 E). Previous work demonstrated how D262A ^2^ and F296A ^4^ mutations in the same binding site in SACK1G disrupt binding to CK1α. Accordingly, BS5 was explored in more detail. The ten ligands grouped together in BS5 are adjacent to the two ligands in BS1 and for simplicity both sites were combined. This site of interest is formed by thirteen residues (Figure 1 F). Figure 1 G displays the relation between the missense enrichment score (MES) on the Y axis and the normalised Shenkin divergence score (*N_Shenkin_*) on the X axis. Positions that are conserved (*N_Shenkin_* < 50) and missense-depleted (MES < 0) are located on the bottom-left quadrant and are the most likely to have a role in the protein’s function since they are constrained both across members of the family and within the human population. Using these two measures of evolutionary constraint, the thirteen residues were ranked by the likelihood of influencing FAM83B function by disturbing CK1α binding (Supplementary Table 1). Residues were stratified in three classes by their divergence and missense-enrichment scores: high likelihood (V149, F273, L233, L179 and D235), medium likelihood (Y277, Y177, D270 and C236) and low likelihood (A151, R274, R206, R208).

**Figure 1.**
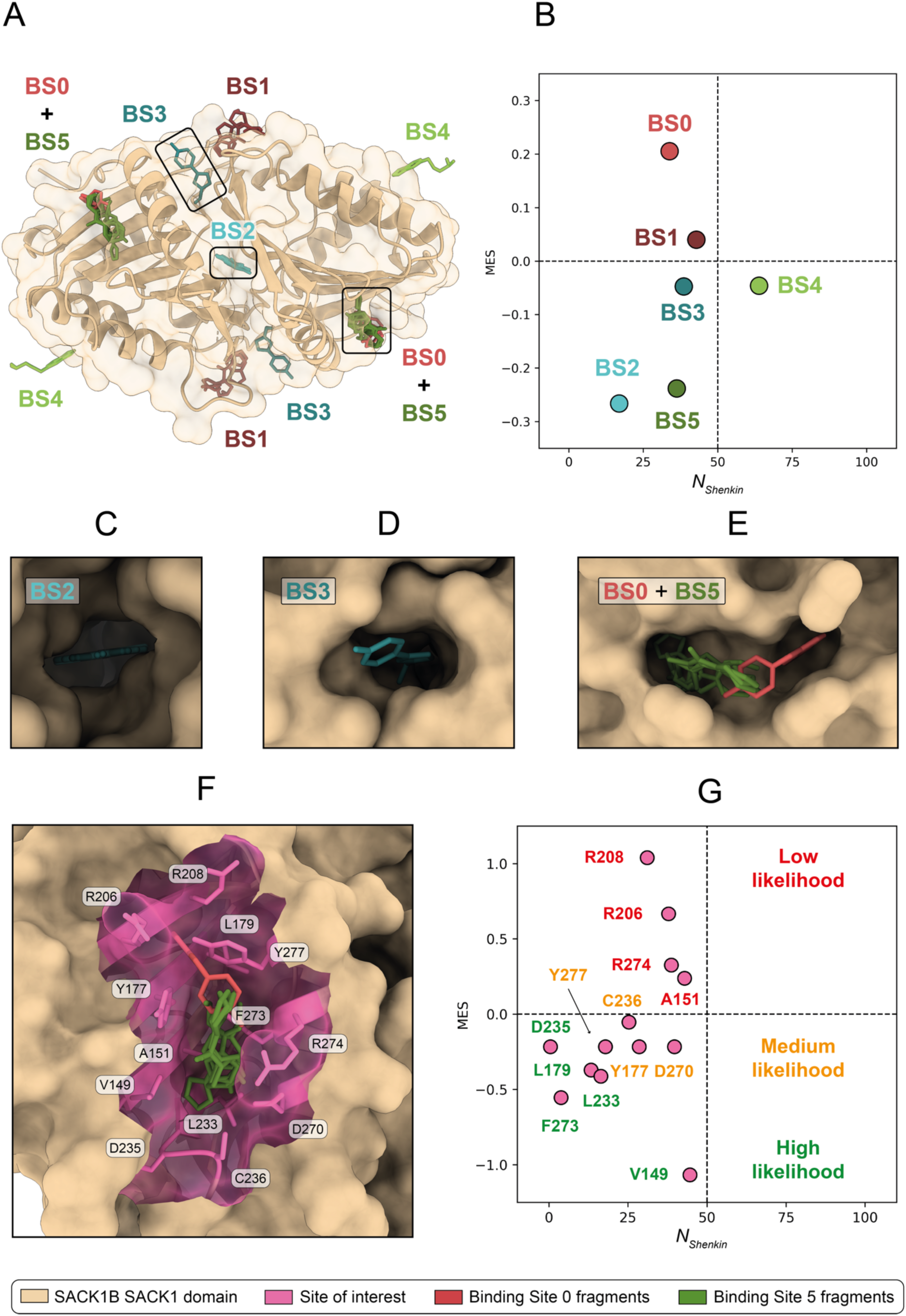
SACK1/FAM83B fragment screening sites. Twenty-four fragments across eleven PDB structures were grouped into six binding sites with the method of Utgés *et al*. ^35^. **(A)** Six defined fragment binding sites for SACK1B/FAM83B on the biologically functional dimer exemplified with PDB: 5QHN ^24^; **(B)** Missense enrichment score (MES) *vs* normalised Shenkin divergence score (*N_Shenkin_*), or *conservation plane*, for the six SACK1B/FAM83B binding sites. Dashed lines indicate conservation (*N_Shenkin_* < 50) and missense depletion (MES < 0) thresholds. Sites on the bottom left quadrant are conserved across homologues, depleted in missense variation and more likely to have a role in the protein’s function; **(C)** Binding Site 2 – PDB: 5QHS ^29^; **(D)** Binding Site 3 – PDB: 5QHL ^22^; **(E)** Binding Site 0 + Binding Site 5 – PDB: 5QHQ ^27^. These two sites are very close to each other and were merged into a single site of interest (SI); **(F)** SACK1B/FAM83B site of interest composed of BS0 and BS5 binding residues; **(G)** Conservation plane for the thirteen residues forming the site of interest. Residues were split into three categories by their potential likelihood of function by their conservation and missense depletion scores. Structure visualisation with ChimeraX ^47^.

### SACK1G dimer modelling

There are no experimentally determined structures for human protein SACK1G. Accordingly, a structural model for the SACK1G SACK1 domain dimer was obtained with AlphaFold Multimer (AFM) (Figure 2) and structurally compared to the monomer model available in AlphaFold DB (AFDB). Both models presented very high confidence (pLDDT > 90) for most of the domain except for a loop at positions 74-128 (Supplementary Figure 1 A-B). This agrees with disorder prediction from MobiDB ^60^. This loop, as well as a confidently predicted *3-helix region* towards the N-terminus at residues 29-73, are the locations where the two models differ. The model quality in this *3-helix region*, as well as its position relative to the structured core of the domain improve when modelling SACK1G as a dimer, indicated by higher pLDDT and lower PAE (Supplementary Figure 2). These helical regions were also predicted in the same location by JPred, lending confidence in their structure. Focusing on the well-structured region of the SACK1 domain (129-315), the models are structurally very similar with RMSD = 0.51 Å (Supplementary Figure 1 C-D). This structural similarity remains when comparing to the AFDB model of SACK1B and an experimental structure 5QHN ^24^ with RMSD = 0.65 Å (Supplementary Figure 1 E-F). This structural similarity suggests that a ligand binding site homologous to the one identified with the fragment screen of SACK1B might exist in SACK1G.

**Figure 2.**
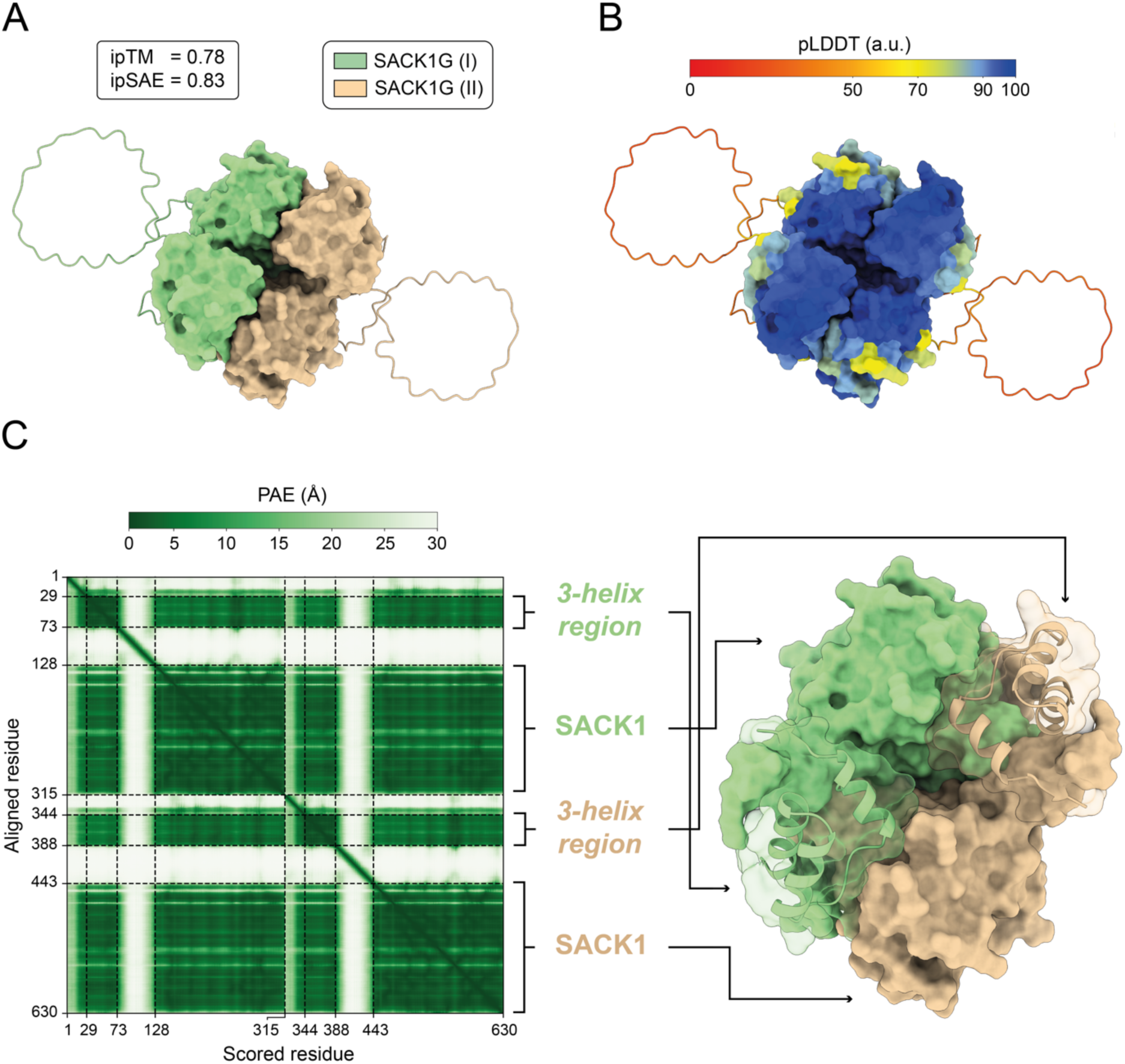
SACK1G/FAM83G SACK1 domain dimer model. A Colabfold implementation of AlphaFold Multimer (AFM) was employed to predict the structure of the SACK1G/FAM83G SACK1 domain dimer, from residues 1-315. ipTM = 0.78 and ipSAE = 0.83. **(A)** Model coloured by chain. Residues with pLDDT > 70 are displayed with a surface. Low-confidence regions are displayed with a ribbon; **(B)** Model coloured by pLDDT; **(C)** PAE matrix for the model. There are two defined low PAE regions or *subdomains*. The core of the globular SACK1 domain (128-315 and 443-630) for the first and second SACK1G/FAM83G copy, and a *3-helix region* at 29-73 and 344-388.

Both the AFM and AFDB models of SACK1G, as well as AFDB model of SACK1B, contain an extra region of structural coverage compared to experimentally resolved structures of SACK1B, e.g., 5QHN. This region includes residues 29-73 in SACK1G which fold into three α-helices (α1, α2 and α3) and is accordingly referred to as *3-helix region* (Figure 3 A). Previous studies identified missense variants SACK1G^A34E 11^ and SACK1G^R52P 13^ as pathogenic leading to the development of palmoplantar keratoderma (PPK) in humans and dogs, respectively. Both variants fall within this 3-helix region suggesting this region is relevant for SACK1G function. A34 is located on α1 and forms a network of hydrophobic interactions with several residues on α2 including F46, V49, L50, I55 and F58. A change to a larger negatively charged residue like glutamate would very likely alter this network of contacts disrupting the helix packing. R52 does not interact with many residues within the region but could play a role in binding to CK1α as suggested by Glennie *et al*. ^9^.

**Figure 3.**
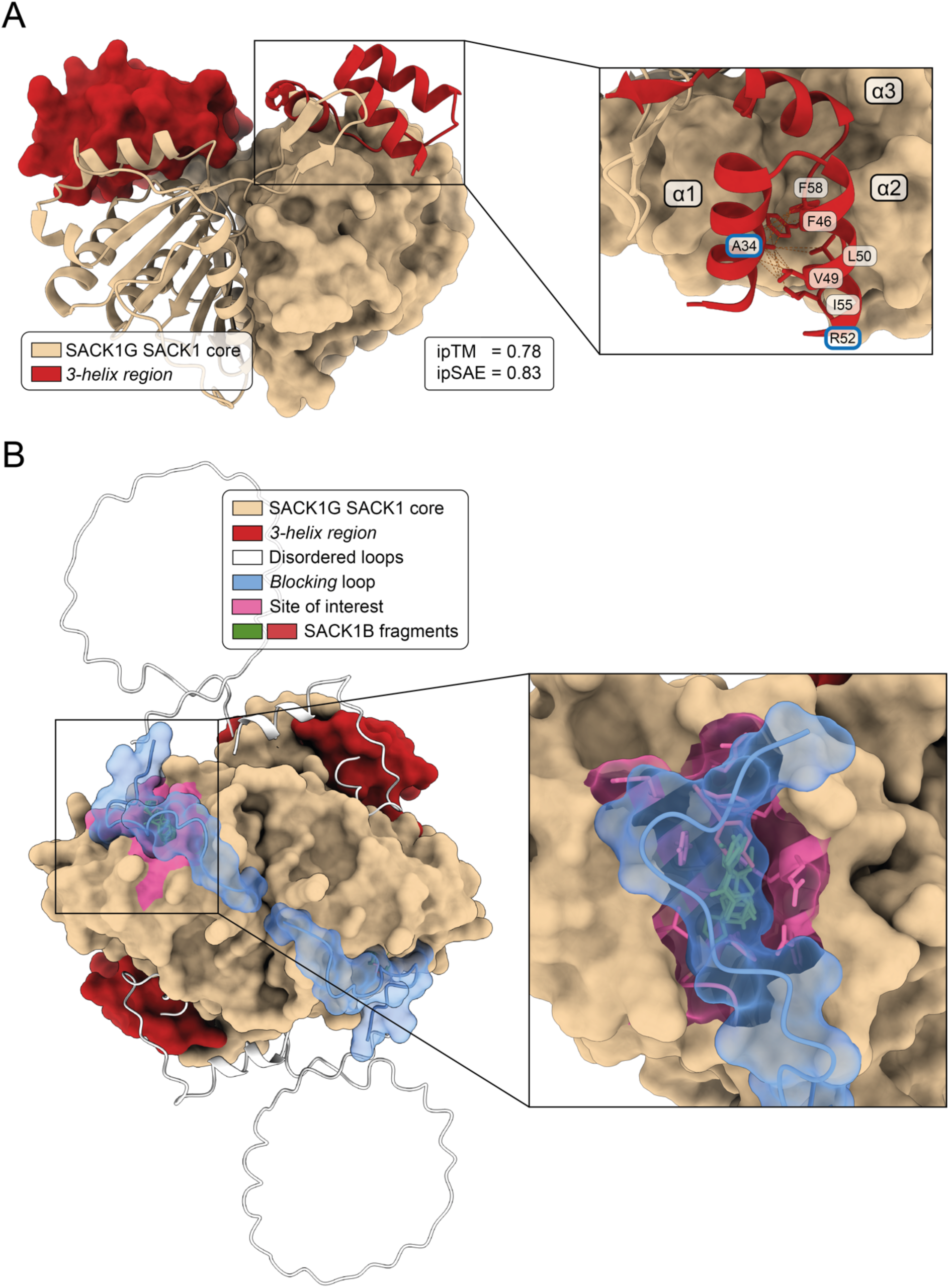
SACK1G/FAM83G dimer model analysis. **(A)** FAM83G dimer model (residues with pLDDT > 70). ipTM = 0.78 and ipSAE = 0.83. The *3-helix region* located at residues 29-73 is highlighted. There are two previously reported pathogenic variants within this region: SACK1G/FAM83G^A34E 11^ and SACK1G/FAM83G^R52P 13^. A34 is predicted to form a network of hydrophobic interactions with five residues. A mutation on this position might disrupt the packing of this region, therefore leading to an abnormal dysfunctional protein product. R52 might play a role in CK1α binding ^9^; **(B)** Full SACK1G/FAM83G dimer model. High-confidence residues (pLDDT > 70) are represented with surface and low-confidence ones with ribbon. Fragments binding to the SACK1B/FAM83B site of interest fit nicely into the homologous pocket in SACK1G/FAM83G. This site of interest seems to be blocked or covered by a confidently predicted loop located at 129-145, which might act as a regulatory element.

Additionally, our SACK1G model included a confidently-predicted loop (pLDDT > 70; PAE < 5 Å) that is not present in the homologous SACK1B structures determined by X-ray crystallography. This loop is found at residues 128-145 and it seems to be covering or blocking the homologous SACK1G fragment binding site of interest defined from SACK1B (Figure 3 B). If this site is indeed functionally relevant, as suggested by our analysis of solvent accessibility, evolutionary divergence and enrichment in variation, it could be that this loop has a protective role and blocks the site to ensure SACK1G function. This loop might be able to undergo a conformational change and move out of the pocket leaving the site accessible for ligand binding and allosteric modulation.

### SACK1G binding site analysis

A binding pocket of interest was defined on SACK1G thanks to the structural similarity between the experimentally determined structures of SACK1B and the AlphaFold Multimer model of the SACK1G dimer (Figure 4 A). Figure 4 B illustrates the evolutionary divergence and enrichment in human missense variation of the thirteen residues forming this site in the same way as it was done for the same pocket in SACK1B (Figure 1 G). Residues were ranked based on likelihood of function using these two scores and stratified in three categories again: high, medium and low likelihood. Those residues presenting lower divergence and higher variant depletion were on the top of the list as they were constrained both across members of the family and within human. Six residues out of the thirteen forming the pocket were selected for alanine scanning and subsequent experimental functional assay. These were: V175, Y204, I206, R235, F260 and Q301. This residue selection, in conjunction with D262 and F296 – which are already known to abolish CK1α binding if mutated to alanine ^2,4^ – includes positions across the different conservation and variation classes defined in Figure 4B and offers good structural coverage of the pocket.

**Figure 4.**
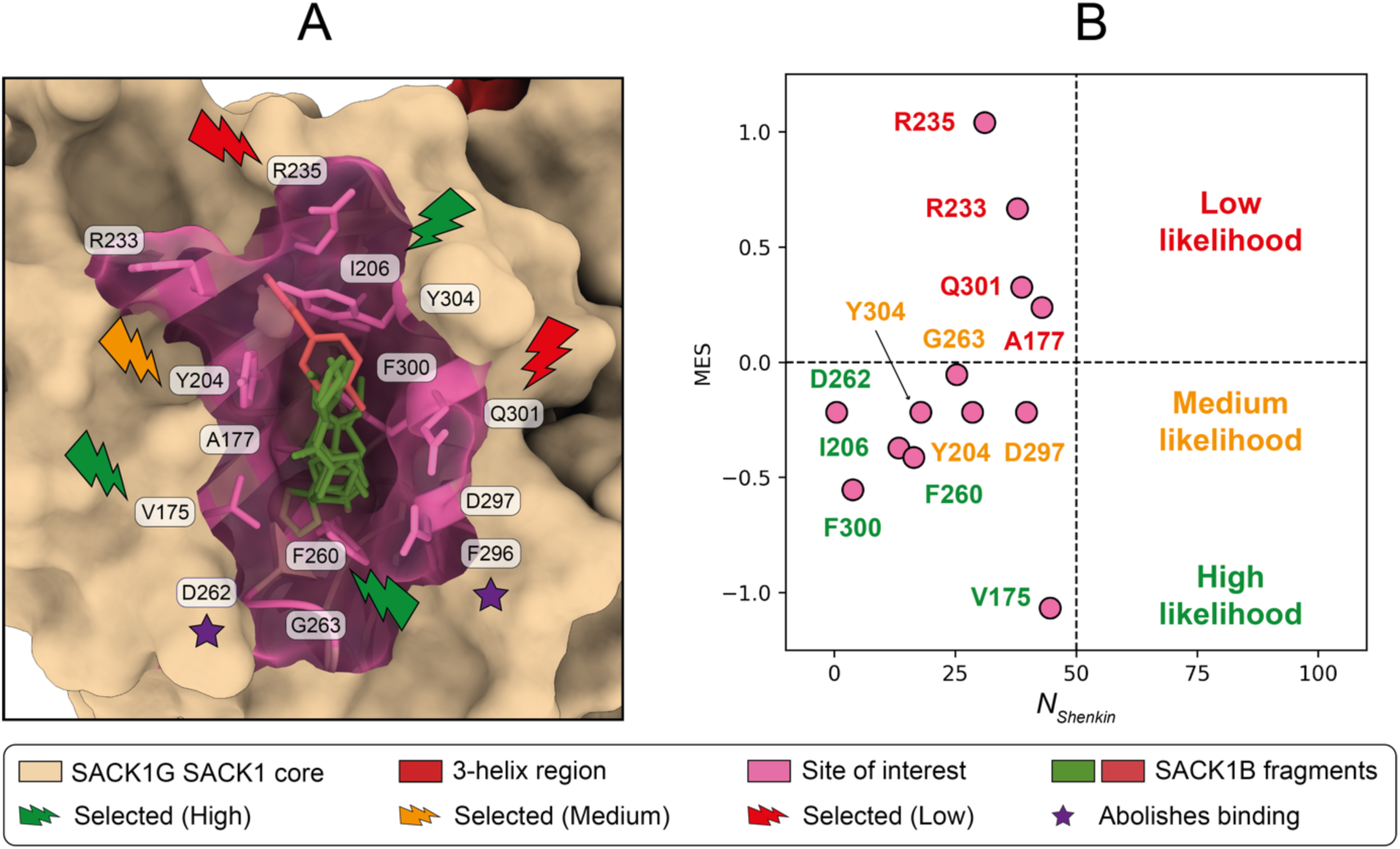
SACK1G/FAM83G binding site analysis. This binding site on SACK1G/FAM83G is defined by the superposition of the SACK1B/FAM83B site of interest to the SACK1G/FAM83G AFM model obtained in this work. **(A)** SACK1G/FAM83G site of interest. Pocket residues are coloured in pink and residues selected for experimental alanine scan, as well as those known to abolish CK1α binding are highlighted; **(B)** Conservation plane for the thirteen residues forming this site in SACK1B/FAM83B. Residues were classified into three groups by their divergence and missense enrichment scores.

### Impact of SACK1G mutants on SACK1G-CK1α binding

SACK1G associates with the Ser/Thr protein kinase CK1α and this association is critical in mediating WNT signalling activation. All four reported PPK pathogenic mutants of SACK1G (A34E, R52P, F58I and R265P) abolish their association with CK1α. Therefore, we sought to decipher how the six chosen mutations, namely V175A, Y204A, I206A, R235A, F260A and Q301A, within the SACK1 domain of SACK1G affect the SACK1G-CK1α interaction in cells. We stably restored the expression of wild type SACK1G, the PPK pathogenic SACK1G^A34E^ mutant or each of the six chosen FAM83G mutants (SACK1G^V175A^, SACK1G^I206A^, SACK1G^Q301A^, SACK1G^F260A^, SACK1G^Y204A^ and SACK1G^R235A^) all with a C-terminal GFP tag in *SACK1G*^-/-^ DLD-1 colorectal cells ^8,9^ (Figure 5A). As expected, anti-GFP IPs from cells expressing wild type SACK1G co-precipitated endogenous CK1α but not from those expressing SACK1G^A34E 5^ (Figure 5B). Anti-GFP IPs from cells expressing SACK1G^V175A^, SACK1G^Q301A^, SACK1G^F260A^ and SACK1G^R235A^ also co-precipitated endogenous CK1α (Figure 5B), indicating that these mutations by themselves do not appear to interfere with the SACK1G-CK1α interaction. Excitingly, however, anti-GFP IPs from cells expressing SACK1G^I206A^ and SACK1G^Y204A^ did not co-precipitate endogenous CK1α (Figure 5B).

**Figure 5.**
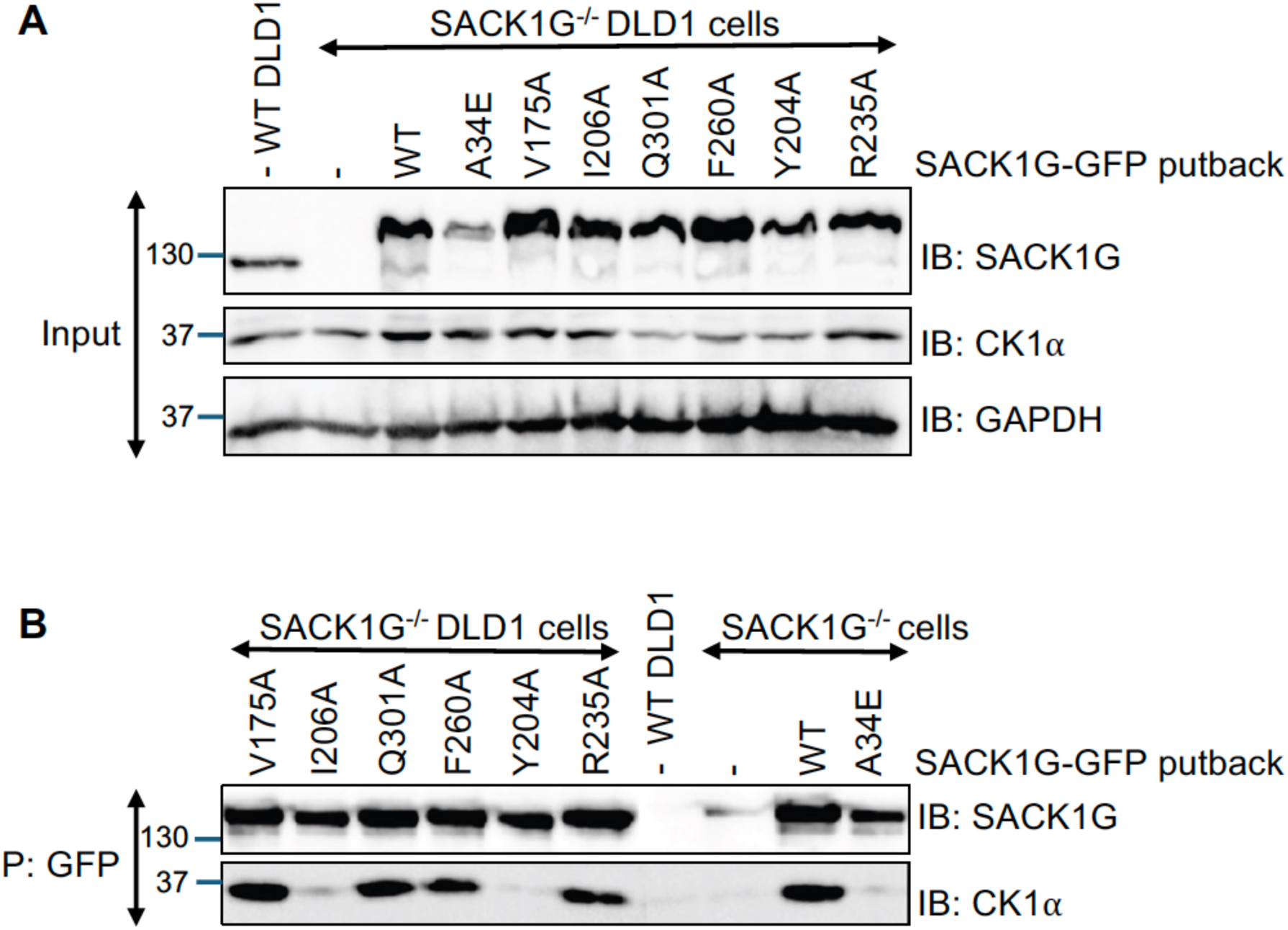
FAM83G/SACK1G I206A and Y204A mutants abolish SACK1G-CK1α binding. **(A)** The indicated variants of SACK1G derived from Figure 4 and Table 1 were stably expressed in SACK1G^-/-^ DLD1 cells with a GFP tag at the C-terminus using retroviruses encoding these mutants. Extracts (20 µg protein) were subjected to SDS-PAGE and samples transferred to PVDF membrane, which was subjected to immunoblotting using the indicated antibodies; **(B)** Cleared extracts (1 mg protein) from (A) were subjected to anti-GFP immunoprecipitation (IP) and IPs subjected to SDS-PAGE, transfer to PVDF membrane and immunoblot analysis using the indicated antibodies.

**Table 1.** SACK1G/FAM83G site of interest. Thirteen residues forming the site of interest on SACK1G/FAM83G. This ligand binding site of interest is homologous to the one defined from the analysis of the fragment screening experiment of SACK1B/FAM83B. Evolutionary divergence is measured with the normalised Shenkin divergence score (*N_Shenkin_*). Enrichment in human missense variation is quantified with the missense enrichment score (MES). Residues were ranked by their conservation and missense scores. The scores are the same as for the SACK1B/FAM83B residues as they are calculated from the multiple sequence alignment column and the residues in both proteins are homologous, i.e., share common ancestry and align in the same column. Group shows the functional likelihood class assigned to each residue based on their ranking. Residues selected for experimental validation are indicated with a check mark (✓).

| Rank | Residue | $N_{Shenkin}$ | MES | Group | Selected |
| --- | --- | --- | --- | --- | --- |
| 1 | V175 | 45 | −1.07 | High likelihood | ✓ |
| 2 | F300 | 4 | −0.55 | High likelihood | X |
| 3 | F260 | 16 | −0.41 | High likelihood | ✓ |
| 4 | I206 | 13 | −0.37 | High likelihood | ✓ |
| 5 | D262 | 1 | −0.22 | High likelihood | X |
| 6 | Y304 | 18 | −0.22 | Medium likelihood | X |
| 7 | Y204 | 29 | −0.22 | Medium likelihood | ✓ |
| 8 | D297 | 40 | −0.22 | Medium likelihood | X |
| 9 | G263 | 25 | −0.05 | Medium likelihood | X |
| 10 | A177 | 43 | 0.24 | Low likelihood | X |
| 11 | Q301 | 39 | 0.33 | Low likelihood | ✓ |
| 12 | R233 | 38 | 0.67 | Low likelihood | X |
| 13 | R235 | 31 | 1.04 | Low likelihood | ✓ |

### Targeted degradation of CK1α co-deplete wild type SACK1G but not SACK1G^I206A^ and SACK1G^Y204A^ mutants

Previously, we reported that lenalidomide degrades the SACK1F-CK1α complex by recruiting CK1α to the CRBN-CUL4A E3 ligase complex for ubiquitin-mediated proteasomal degradation ^7^. Recently, two derivatives of lenalidomide, DEG-77 ^61^ and SJ3149 ^62^, were shown to be potent degraders of CK1α acting via the CRBN-CUL4A E3 ligase complex. Using DEG-77 and SJ3149, we were able to demonstrate a robust co-depletion of the SACK1G-CK1α, as well as the SACK1F-CK1α complex ^63^. We postulated that SACK1G^I206A^ and SACK1G^Y204A^ mutants would be resistant to degradation by DEG-77 as they no longer interact with CK1α. First, we optimised DEG-77 treatment conditions for co-depletion of SACK1G-CK1α complex in DLD-1 cells. Treatment of DLD-1 cells with increasing doses of DEG-77 for 24 h led to a dose-dependent depletion in levels of CK1α, SACK1G and SACK1F compared to DMSO treatment, with maximal depletion observed at concentrations above 100 nM (Figure 6A). When cells were treated with 100 nM DEG-77 over a course of 24 h, co-depletion of CK1α, SACK1G and SACK1F was observed after 3 h, with maximal depletion observed at 24 h (Figure 6A-B). Next, *SACK1G*^-/-^ DLD-1 cells stably expressing WT SACK1G-GFP, SACK1G^A34E^-GFP, SACK1G^I206A^-GFP and SACK1G^Y204A^-GFP were treated with 100 nM DEG-77 for 24 h (Figure 6C). In cells expressing WT SACK1G-GFP, compared to DMSO control, DEG-77 robustly depleted both CK1α and SACK1G-GFP. In contrast, in cells expressing SACK1G^A34E^-GFP, SACK1G^I206A^-GFP and SACK1G^Y204A^-GFP, while CK1α was robustly degraded, no co-depletion of SACK1G^A34E^-GFP, SACK1G^I206A^-GFP and SACK1G^Y204A^-GFP was observed (Figure 6C), confirming that these mutants abolish their interaction with CK1α. Further analysis through multiple sequence alignment (Supplementary Figure 3) revealed that Y204 is conserved across all SACK1 members, except for SACK1A, which interestingly exhibits the weakest binding affinity to CK1α ^2^. I206 is conserved in SACK1F and SACK1G, while other members have a similar hydrophobic leucine residue at this position.

**Figure 6.**
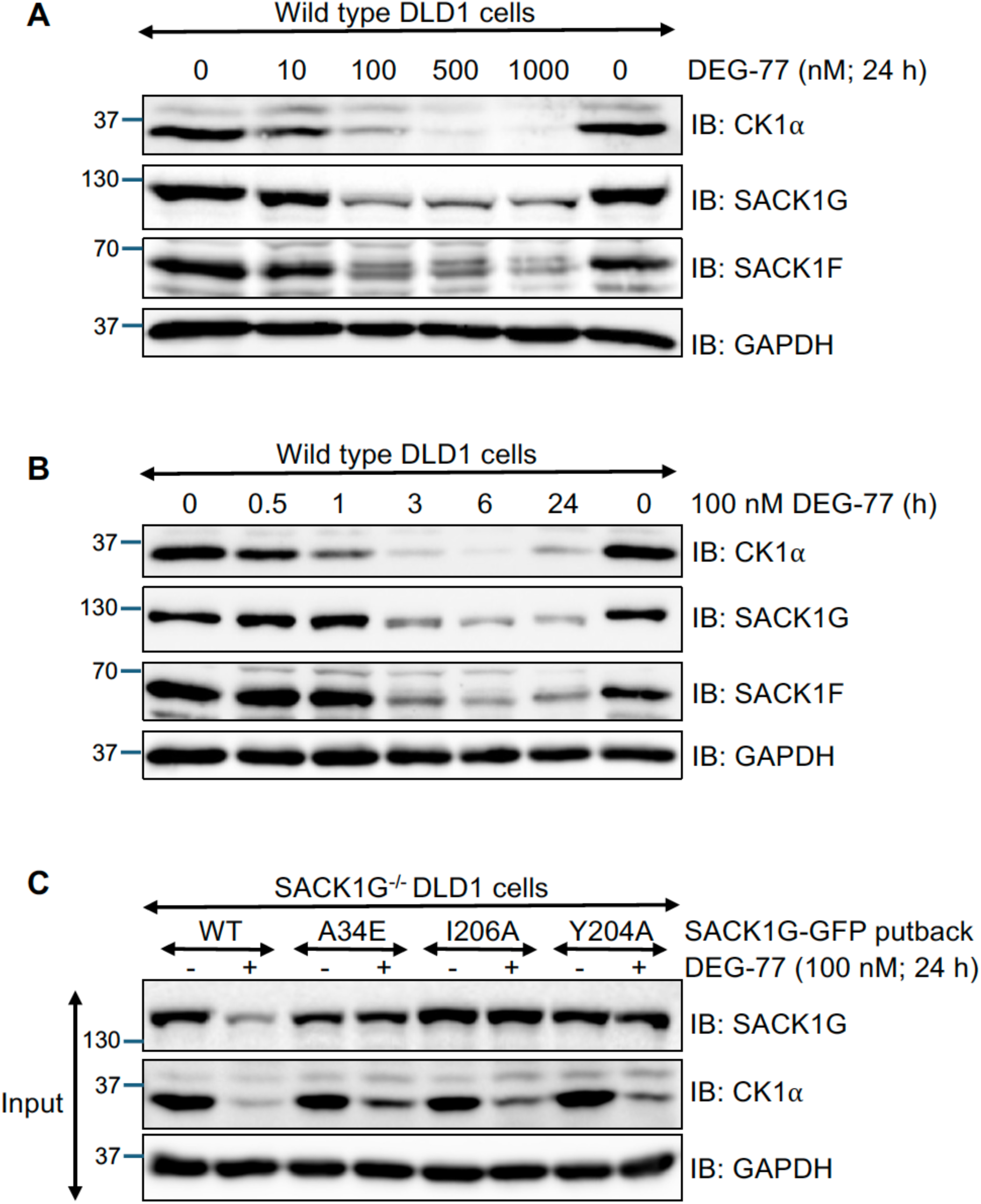
DEG-77 fails to degrade SACK1G/FAM83G I206A and Y204A mutants. **(A&B)** Wild type DLD1 cells were subjected to treatment with **(A)** the indicated concentrations of DEG-77 for 24 h and **(B)** 100 nM DEG-77 for the indicated times prior to lysis. Extracts (20 µg protein) were subjected to SDS-PAGE and samples transferred to PVDF membrane, which was subjected to immunoblotting using the indicated antibodies; **(C)** Wild type (WT) SACK1G-GFP or the indicated mutants of SACK1G-GFP were stably expressed in SACK1G^-/-^ DLD1 cells using retroviruses encoding these mutants. Cells were treated with 100 nM DEG-77 for 24 h prior to lysis. Extracts (20 µg protein) were subjected to SDS-PAGE and samples transferred to PVDF membrane, which was subjected to immunoblotting using the indicated antibodies.

## Discussion

Based on high-throughput fragment screening data, we previously reported a method to define ligand-binding clusters on proteins to ascertain impact on protein function. In this manuscript we focused on one protein, namely SACK1B, and identified a ligand-binding pocket within its SACK1 domain as potentially functionally significant. There are some reports suggesting the role of SACK1B in promoting cancer cell proliferation and mediating resistance to TKI inhibitors ^64,65^, however its precise molecular role remains elusive. Because of this, we modelled the structure of the conserved SACK1 domain of the better characterised SACK1G. We classified residues near the ligand-binding site into high, medium and low likelihood of functional impact based on conservation and missense depletion scores. We then mutated two residues from each category from SACK1G to test their effect on CK1α binding. Excitingly, two mutations, namely I206A (high likelihood) and Y204A (medium likelihood), completely abolished the binding to CK1α in cells. Consistent with this loss of interaction with CK1α, degraders of CK1α that also co-degrade native SACK1G failed to degrade I206A and Y204A mutants. Loss of CK1α binding and subsequent inhibition of WNT signalling underlies the pathogenesis of PPK caused by *SACK1G* mutations within the SACK1 domain ^5,9^. It would be interesting to investigate whether I206 and Y204 mutations are also associated with PPK or if loss of SACK1G-CK1α interaction by I206A and Y204A mutations also lead to inhibition of WNT signalling ^1,4,5,9^. Within the vicinity of this predicted ligand-binding pocket on SACK1G are conserved residues D262 and F296, which have previously been shown to play critical role in CK1α binding and activation of WNT signalling ^4^. Therefore, developing the current low-affinity small fragments into selective ligands within this pocket may yield ligands with potential functional consequence that might prove useful tools for exploring the cellular functions of SACK1G-CK1α complexes. Unlike SACK1G which selectively binds CK1α, SACK1B binds to CK1α, CK1δ and CK1ε ^2^. Therefore, it would be interesting to test whether the equivalent mutations on FAM83B, namely L179A and Y177A, also abolish interactions with CK1α, CK1δ and CK1ε.

## Conclusions

The conclusions resulting from this work are as follows:

- Modelling SACK1G as its biologically functional dimer improves the quality of the structure prediction relative to that of a monomeric model.
- A C1 site in SACK1G – as predicted by Utgés *et al.* ^35^ – is experimentally shown to be functional, as mutations within the site abolish binding to its partner CK1α.

## Statistics and reproducibility

PDB accession codes for the eleven SACK1B/FAM83B fragment screening structures are the following: 5QHI ^19^, 5QHJ ^20^, 5QHK ^21^, 5QHL ^22^, 5QHM ^23^, 5QHN ^24^, 5QHO ^25^, 5QHP ^26^, 5QHQ ^27^, 5QHR ^28^ and 5QHS ^29^. Colabfold v1.5.2 was utilised for 3D structure prediction. Computational analysis was carried out primarily with the following Python libraries: Matplotlib ^66^, Pandas ^67,68^ and SciPy ^69^ and Seaborn ^70^.

## Data availability

All the 3D structure models and predictions analysed in this work can be found in the following GitHub repository: https://github.com/bartongroup/SACK1G-analysis ^71^. The raw data generated for this study is available at Mendeley (link will be added here following revisions).

## Code availability

The code developed to carry out this analysis is available from our GitHub repository: https://github.com/bartongroup/SACK1G-analysis.

## Competing interests

The Sapkota laboratory receives or has received sponsored research support from Amgen, Boehringer Ingelheim, GlaxoSmithKline and Johnson & Johnson. The authors declare no other competing interests.

## Funding

This work was supported by grants to G.J.B. from UKRI-Biotechnology and Biological Sciences [BB/X018628/1] and Wellcome Trust [101651/Z/13/Z; 218259/Z/19/Z] and by grants to G.P.S. [UKRI Medical Research Council: MC_UU_00018/6 and MC_UU_00038/6]. G.P.S. is also supported by Boehringer-Ingelheim (through the Division of Signal Transduction Therapy). J.S.U. was supported by a BBSRC EASTBIO Ph.D. Studentship [BB/J01446X/1]. L.G. and B.L.C. are supported by the MRC PPU PhD studentships.

## Author contributions

J.S.U. and G.J.B. conceived, designed and developed the computational side of the work. D.L.Z.X., B.L.C., and L.G. performed biochemical and cell-based experiments, analysed data and compiled figures. J.S.U. developed the code and performed the bioinformatics analysis.

T.J.M. and N.T.W. generated constructs used in this study. J.S.U., G.J.B, and G.P.S. wrote, reviewed and edited the manuscript. G.P.S. and G.J.B. secured funding and supervised the project.

## Acknowledgements

We thank Dr James Abbot for the Colabfold implementation on the cluster and for advice on how to run predictions. We also thank the IT service of the University of Dundee for their support of the HPC infrastructure this study was carried out on. We thank the members of G.P.S. lab for experimental advice and discussions during the study. We thank the MRC tissue culture facility staff (E. Allen, A. Muir, S. Dalglish, E. Webster and J. Stark), the staff at the DNA Sequencing services (School of Life Sciences, University of Dundee), the cloning, antibody, inhibitor and protein production teams within the MRC PPU Reagents & Services (University of Dundee) coordinated by J. Hastie and the staff at FACS service for their technical and reagents support. We thank Christina Woo (Broad Institute of MIT and Harvard) for providing us with DEG-77.

---

**Supplementary Figure 1.**
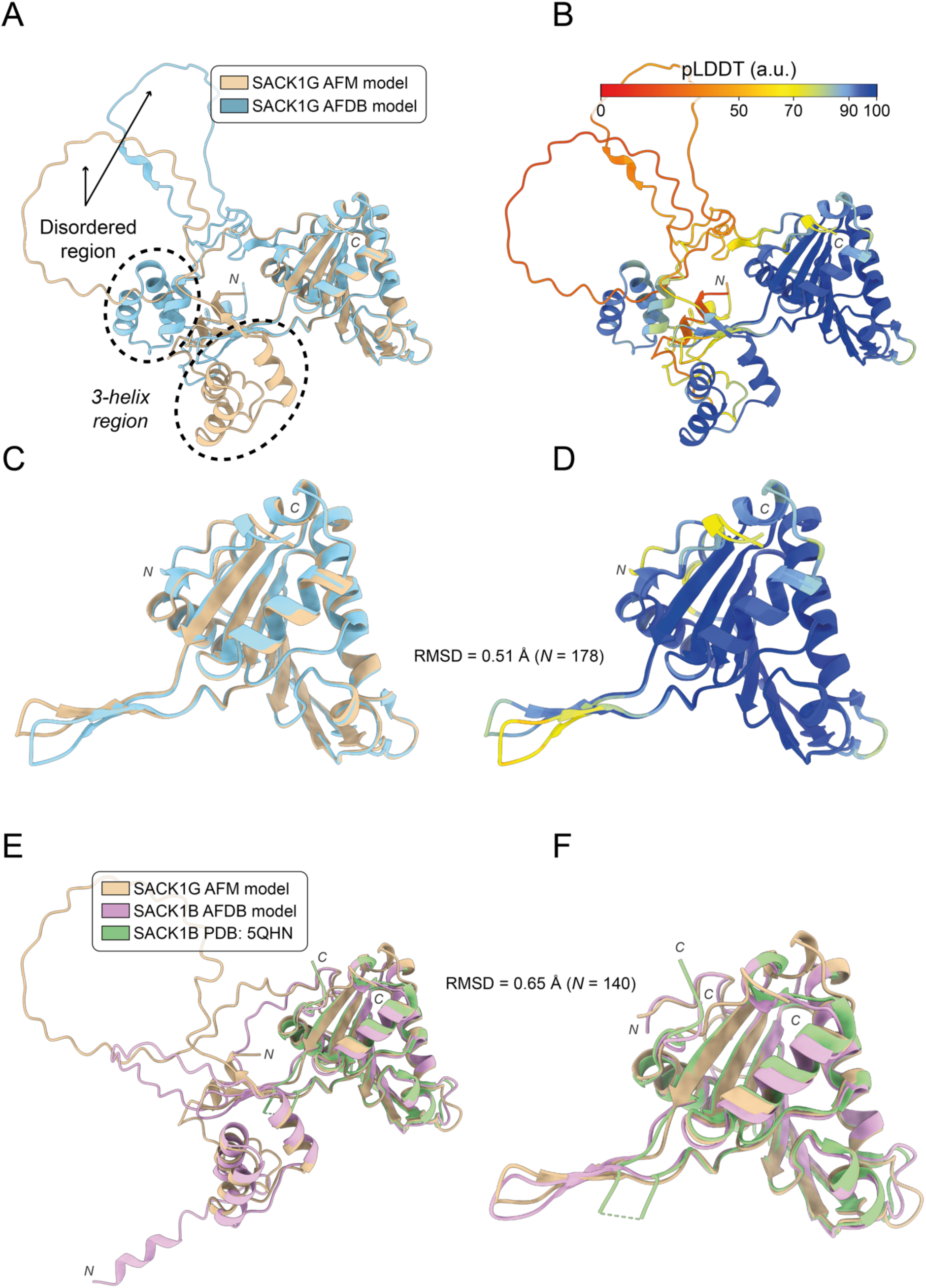
Comparison of SACK1G/FAM83G AlphaFold Multimer and AlphaFold DB models. Our AlphaFold Multimer (AFM) model is a dimer (ipTM = 0.78 and ipSAE = 0.83). The AlphaFold DB (AFDB) is of a monomer. A single copy is included in this figure for simplicity. **(A)** Superposed AFM and AFDB SACK1G/FAM83G models. A novel *3-helix region* – not present in the crystal structures of close homolog SACK1B/FAM83B – is highlighted (29-73) as well as disordered loop region (73-128); **(B)** Models coloured by pLDDT; **(C)** Superposition of SACK1 domain core (129-315) between AFM and AFDB models. These are highly similar, with an RMSD = 0.51 Å across the 178 best aligning residues; **(D)** Superposition of SACK1 domain core models coloured by pLDDT; **(E)** Superposition of our SACK1G/FAM83G model, the FAM83B AFDB model and a FAM83B crystal structure: 5QHN ^24^. The SACK1G/FAM83G model obtained in this work is structurally very similar to the SACK1B/FAM83B AFDB model and crystal structure: RMSD = 0.65 Å; **(F)** SACK1 domain core across our SACK1G/FAM83G model, the SACK1B/FAM83B AFDB model and a SACK1B/FAM83B crystal structure: 5QHN. Structure visualisation with ChimeraX ^47^.

**Supplementary Figure 2.**
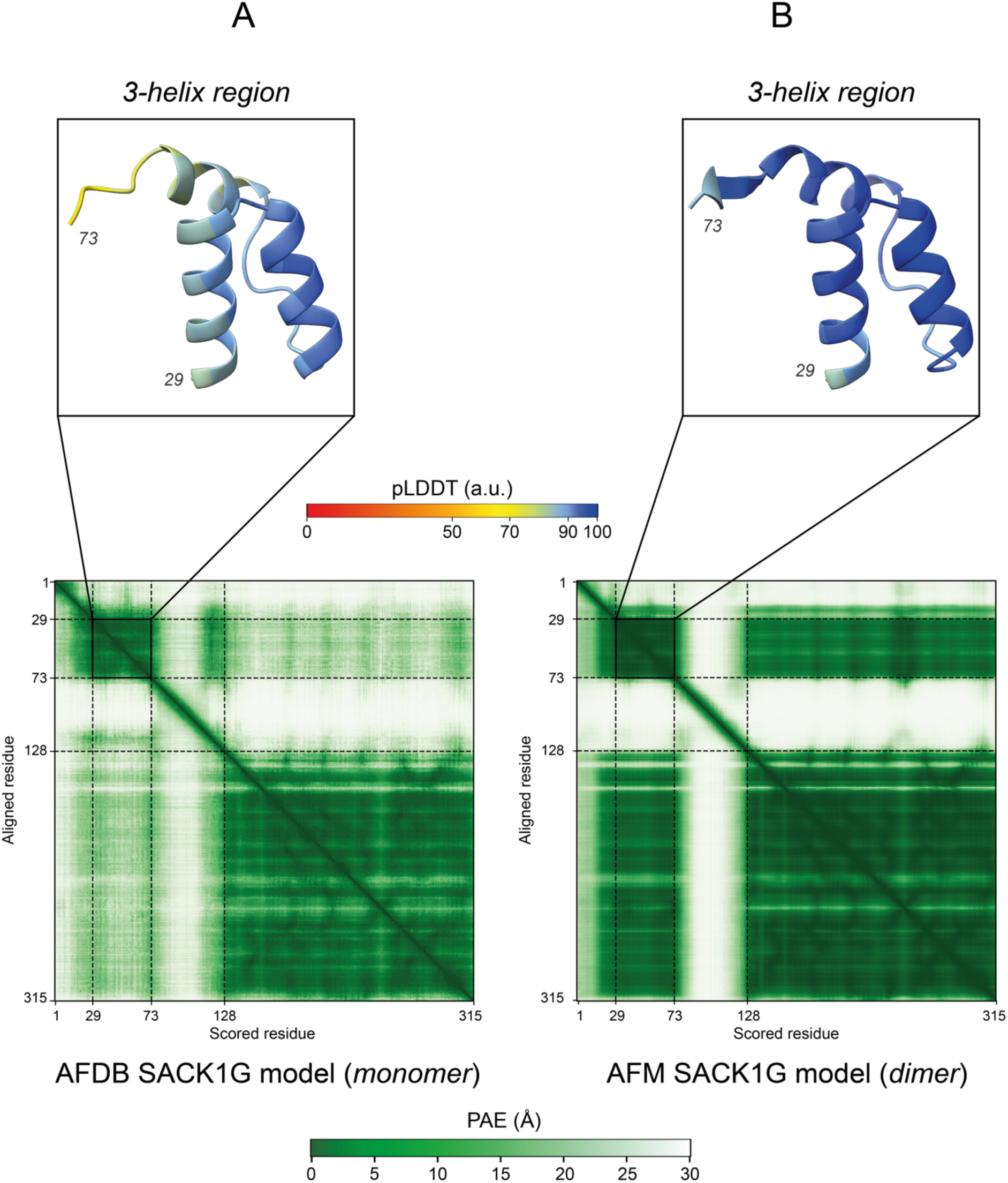
Comparison of the *3-helix region* between SACK1G/FAM83G AFM and AFDB models. Our model – obtained with AFM – is a dimer, and the one from AFDB is of a monomer. Modelling SACK1G/FAM83G as a dimer – which corresponds to its biological assembly – improves the overall confidence of the model in terms of relative orientation between different amino acids. This can be observed by the lower predicted alignment error (PAE), i.e., darker green colour, on the two PAE blocks around the diagonal, which correspond to the *3-helix region* (29-73) and the core of the SACK1 domain (129-315). Additionally, the dimer model is much more reliable regarding the position of these two blocks relative to each other, i.e., the location in 3D space of the *3-helix region* relative to the SACK1 domain core. An improvement on the per-residue confidence (pLDDT) on the *3-helix region* is also observed (darker blue colour).

**Supplementary Figure 3.**
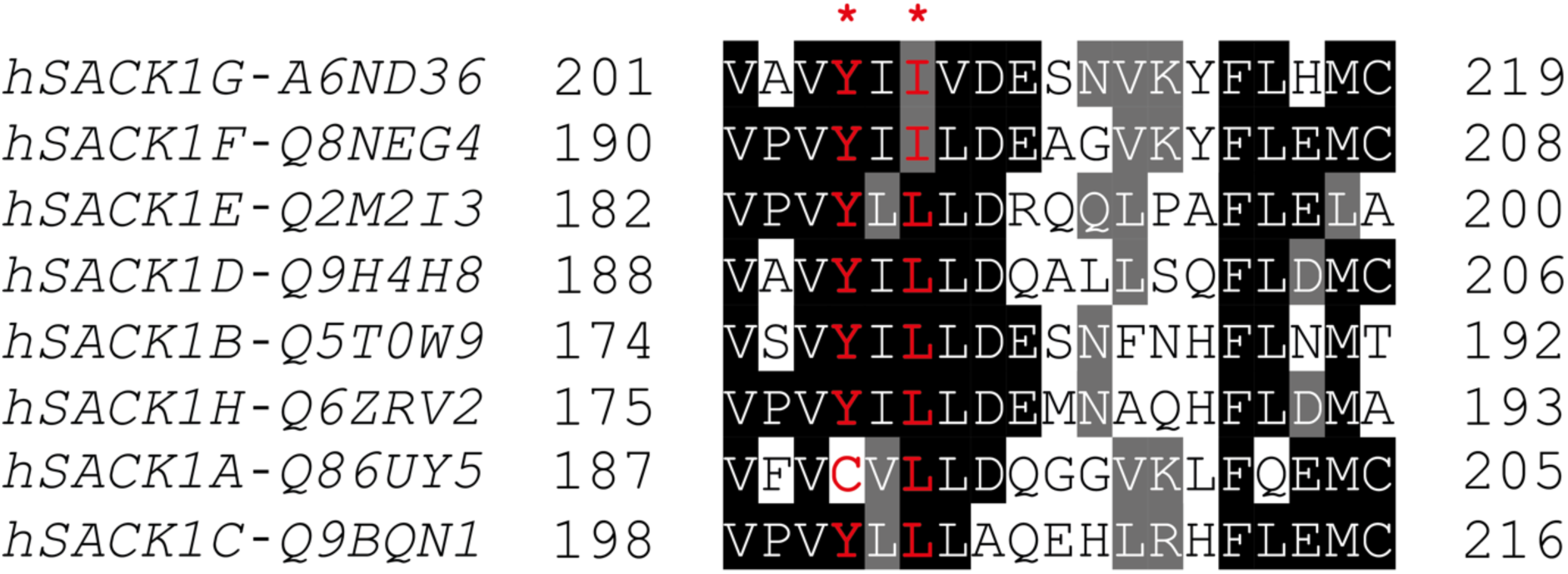
Human SACK1A-H amino acid sequence alignment surrounding resides Y204 and I206 of SACK1G. Sequence alignment was performed using Clustal Omega ^72^ and diagram generated with Jalview ^73,74^. Residues identical in four or more members are shaded in black, residues with similar physicochemical properties in grey. Y204 and I206 of SACK1G and the corresponding residues in other SACK1 proteins are indicated in red font and marked with a star (*).

**Supplementary Table 1.** FAM83B site of interest. Thirteen residues forming the site of interest on SACK1B/FAM83B. This site includes residues from binding sites 0 (BS0) and 5 (BS5) defined from the analysis of the fragment screening experiment of SACK1B. Evolutionary divergence is measured with the normalised Shenkin divergence score (*N_Shenkin_*). Enrichment in human missense variation is quantified with the missense enrichment score (MES). Residues were ranked by their conservation and missense scores. Group shows the functional likelihood class assigned to each residue based on their ranking.

| Rank | Residue | $N_{Shenkin}$ | MES | Group |
| --- | --- | --- | --- | --- |
| 1 | V149 | 45 | −1.07 | High likelihood |
| 2 | F273 | 4 | −0.55 | High likelihood |
| 3 | L233 | 16 | −0.41 | High likelihood |
| 4 | L179 | 13 | −0.37 | High likelihood |
| 5 | D235 | 1 | −0.22 | High likelihood |
| 6 | Y277 | 18 | −0.22 | Medium likelihood |
| 7 | Y177 | 29 | −0.22 | Medium likelihood |
| 8 | D270 | 40 | −0.22 | Medium likelihood |
| 9 | C236 | 25 | −0.05 | Medium likelihood |
| 10 | A151 | 43 | 0.24 | Low likelihood |
| 11 | R274 | 39 | 0.33 | Low likelihood |
| 12 | R206 | 38 | 0.67 | Low likelihood |
| 13 | R208 | 31 | 1.04 | Low likelihood |

